# Insect COI barcoding data as an untapped resource for surveying *Wolbachia* symbioses

**DOI:** 10.64898/2026.06.24.734267

**Authors:** Karol H. Nowak, Mateusz Buczek, Marzena Marszałek, Monika Prus-Frankowska, Catalina Valdivia, Junchen Deng, J. Dylan Shropshire, Piotr Łukasik

## Abstract

1. DNA barcoding of the mitochondrial cytochrome c oxidase I (COI) gene is widely used to characterise insect diversity and distributions; however, its potential to reveal information on species interactions, including host–symbiont associations, remains largely unexplored. Here, we assess whether COI amplicon data can be used to identify *Wolbachia* - one of the most widely distributed bacterial symbionts known to profoundly affect their hosts’ biology.
2. We demonstrate that several commonly used invertebrate COI primer sets perfectly match many reference *Wolbachia* genomes, leading to frequent co-amplification.
3. By screening 7,901 individual-insect COI amplicon libraries obtained with the popular BF3-BR2 primer set, we detected *Wolbachia* sequences in over 35% of samples, revealing that co-amplification is indeed widespread. After removing low-abundance reads, *Wolbachia* detection based on COI amplicons showed over 90% agreement with simultaneously generated 16S-V4 rRNA amplicon data from the same specimens. The degree of agreement, however, varied depending on the thresholds used, among datasets and insect clades.
4. Further, we show that *Wolbachia* abundance inferred from COI amplicons correlated with their abundance in metagenomic datasets for 152 specimens, supporting the quantitative relevance of the signal.
5. Finally, we find that *Wolbachia* COI sequences provide greater phylogenetic resolution than 16S-V4 rRNA data (mean pairwise genetic distance of COI sequences - 9.6%, 16S-V4 rRNA - 2.8%), and the reconstructed *Wolbachia* COI-based genotype network largely agrees with genome-based phylogenies.
6. Collectively, our results demonstrate that off-target *Wolbachia* sequences recovered from standard insect COI barcoding data may reliably detect symbiont presence, provide phylogenetic insight, and guide sample selection for metagenomics. Given the rapid expansion of global insect barcoding initiatives, these findings highlight an opportunity for cost-effective monitoring of their most prevalent bacterial symbionts, offering new perspectives on how host–microbe interactions may shape insect communities.

## Introduction

A series of reports on the global diversity and biomass declines in insects (Wagner et al., 2021) has revealed the need for deeper comprehension of the state of affairs and prompted high-profile monitoring efforts that combine broad collection and creative implementation of high-throughput sequencing-based approaches (Buchner et al., 2025; Chua et al., 2023; Miraldo et al., 2025). In particular, the sequencing of mitochondrial cytochrome c oxidase I (COI) gene, now a standard marker for species identification, has become a popular approach for identifying individual specimens and bulk community samples (Hebert et al., 2003). This approach, widely known as DNA barcoding, relies on short, standardised COI fragments that can be matched against curated reference databases for rapid taxonomic assignment. Indeed, large-scale barcoding initiatives, combined with the development of extensive reference databases, have demonstrated substantial power to reconstruct insect community composition across spatial and temporal gradients (Hartop et al., 2024; Hebert et al., 2025, p. 1). In contrast, the characterisation of biological interactions that underpin community functioning, including predator–prey, host–parasite, and host–symbiont relationships, has lagged behind the expanding species inventories. Investigating these interactions has traditionally required dedicated methodological approaches and has therefore been limited to much smaller spatial and taxonomic scales (Abdala-Roberts et al., 2019).

Coincidentally, the DNA extracted from an insect and the COI amplicon data generated using that DNA can contain information not only about the focal individual but also about interacting organisms, including parasites, parasitoids, dietary items, and microbial associates (Kolasa et al., 2023; Song et al., 2008). A particularly compelling example is *Wolbachia* (Smith et al., 2012). This diverse Alphaproteobacteria genus, comprising at least seventeen well-established supergroups, is considered the most widespread invertebrate symbiont, present in approximately half of all insect species as well as numerous arachnids and nematodes (Kaur et al., 2021). These maternally inherited intracellular bacteria, best known for their ability to alter reproduction, including male killing, feminisation, induction of parthenogenesis, and cytoplasmic incompatibility (Kaur et al., 2021; Shropshire et al., 2020), can also confer protection against natural enemies, especially viruses (Cogni et al., 2021; Teixeira et al., 2008), and contribute to nutrition (Hosokawa et al., 2010; Ju et al., 2020). These combined effects can dramatically alter host biology and fitness, population dynamics and, ultimately, community structure and its dynamics (Łukasik & Kolasa, 2024). Moreover, occasional horizontal transmission enables *Wolbachia* to spread across species and introduce novel phenotypes to different host lineages, influencing their evolutionary trajectories and interaction networks (Shropshire et al., 2026). Overall, there is little doubt that these common bacteria play critically important roles in insect communities. However, the information about their diversity, distribution, and symbiosis dynamics across species remains highly incomplete and largely restricted to model clades including some *Drosophila* and mosquito species (Detcharoen et al., 2026; Scholz et al., 2020). Additionally, *Wolbachia* prevalence, transmission, and phenotypic effects are strongly influenced by environmental conditions, such as temperature (Hague et al., 2022; Ross et al., 2019), making broad spatial and temporal sampling essential for understanding the ecology and dynamics of these symbioses. There is an urgent need to fill this gap and understand *Wolbachia* symbioses across insect taxonomic diversity.

Notably, *Wolbachia* harbours a copy of the cytochrome oxidase I gene - conventionally labelled in bacteria as either *coxA* or *ctaD*, but further referred to as COI, for consistency with insect sequences. Insect and *Wolbachia* gene sequences share many conserved sequence motifs. Consequently, *Wolbachia* COI fragments are often co-amplified by standard invertebrate barcoding primers in PCR reactions (Jalali et al., 2015; Smith et al., 2012). When the resulting PCR product is sequenced using the Sanger method, the insect sequence quality may be negatively affected, or even masked, by the co-amplified *Wolbachia* sequence. Hence, the bacterial co-amplification has been regarded as a nuisance and actively combated, with some primer sets even explicitly designed to minimise co-amplification of *Wolbachia* (Bleidorn & Henze, 2021; Linares et al., 2009) and other Rickettsiales (Park & Poulin, 2020). In other cases, *Wolbachia*-derived sequences have been misidentified as insect barcodes, introducing errors into reference databases (Smith et al., 2012). In the era of high-throughput COI sequencing, *Wolbachia* co-amplification results in distinct bacterial reads that accompany insect reads and may complicate consensus sequence reconstruction, but are easier to detect and filter (Cruaud et al., 2017; Nowak et al., 2025). Here, we ask whether *Wolbachia* co-amplification - long-recognised nuisance - can be reframed as an opportunity: a chance to cost-effectively assess the distribution and diversity of this ecologically relevant symbiont. If this is confirmed and validated, and the limitations of this approach are understood (Stahlhut et al., 2012), insect COI amplicon datasets could become an untapped resource for detecting and characterising the symbioses of this globally distributed microbe.

Our objective was to test whether reliable and biologically meaningful information on *Wolbachia* presence, abundance, and diversity can be extracted from standard insect COI amplicon data. Specifically, we aimed to: (1) assess the extent to which widely used insect barcoding primers match diverse *Wolbachia* reference sequences; (2) characterise the prevalence and relative abundance of *Wolbachia*-derived reads across large insect COI datasets; (3) evaluate the reliability of COI-based *Wolbachia* presence inference by comparison with amplicon data for the bacterial 16S-V4 rRNA marker region; (4) test the concordance between *Wolbachia* abundance estimates based on COI amplicon and metagenomic data; and (5) compare the phylogenetic resolution provided by *Wolbachia* COI sequences, 16S-V4 rRNA sequences and genome-level data. To address these aims, we conducted systematic comparisons of frequently used COI primers against *Wolbachia* reference sequences, followed by large-scale analyses of paired COI and 16S rRNA amplicon data for 7,901 insect specimens. We further integrated metagenomic data from 152 specimens to benchmark abundance estimates and phylogenetic inference. Through this integrative framework, we evaluate whether routine insect barcoding data can be repurposed to illuminate host–symbiont associations at a scale commensurate with contemporary biodiversity monitoring.

## Methods

### In silico assessment of primer specificity

To assess the extent to which commonly used insect metabarcoding COI primers may co-amplify *Wolbachia* sequences, a reference set of *Wolbachia* sequences was obtained: all complete *Wolbachia* genomes available in NCBI, as well as contig- or scaffold-level assemblies not assigned to supergroups A or B based on the five multilocus sequence typing (MLST) loci (downloaded January 2023), were retrieved. Genome annotation was performed using Prokka v1.14.5 (Seemann, 2014), and COI gene sequences were subsequently extracted from the annotated genomes.

The extracted sequences were aligned using MAFFT v7.515 (Katoh & Standley, 2013) and manually curated. Alignments were trimmed to the 709 bp around the Folmer region of COI. Five sequences not covering the full target region were discarded. The remaining 185 sequences were dereplicated, resulting in 73 unique COI sequences representing thirteen *Wolbachia* supergroups (Supplementary Table S1). A phylogenetic ordering of these sequences was inferred using IQ-TREE2 (Minh et al., 2020), with *Anaplasma centrale* (CP001759) used as an outgroup.

*Wolbachia* consensus sequences were generated by assigning IUPAC ambiguity codes for positions where alternative nucleotides were present with at least 10% frequency, separately for each supergroup. A metaconsensus sequence was obtained similarly, however, with each *Wolbachia* supergroup weighted equally to account for overrepresentation of supergroups A and B. The insect consensus sequence was reproduced using data from Elbrecht et al., 2019.

A set of 57 COI primers (28 forward and 28 reverse) commonly used in arthropod barcoding and metabarcoding was compiled from two comprehensive publications (Elbrecht et al., 2019; Tournayre et al., 2024). (Supplementary Table S2). These primers target or flank the 658 bp Folmer region. Additionally, a *Wolbachia*-specific forward primer used in MLST analyses (Baldo et al., 2006) was included. The corresponding reverse *Wolbachia* primer did not overlap with the target region and was therefore ignored.

To evaluate primer–template compatibility, each primer was aligned against the *Wolbachia* COI reference sequences. Nucleotide-level mismatch rates were calculated as the proportion of dereplicated reference sequences containing a primer-incompatible nucleotide. Degenerate positions were treated as matches when at least one nucleotide variant was compatible with the reference base. The primer-level mismatch was defined as the proportion of reference sequences with at least one nucleotide mismatch across the full primer length.

To assess the amplification breadth of COI primers among bacteria other than *Wolbachia*, we have extracted all copies of genes annotated as "aa3-like cytochrome oxidase I" (*ctaD* or *coxA*) from all 59,318 bacterial RefSeq genomes available at NCBI on 18th April 2026, alongside their taxonomy classifications. We then checked which gene copies are perfectly matched by the COI primers listed in the Supplementary Table S2. For each of the 2,348 bacterial genera represented, the proportion of genomes encoding at least one *coxA*/*ctaD* copy was determined. The presence of exact matches, as well as matches compatible with degenerate nucleotide positions in the primers, was then evaluated across these gene copies.

### Sample collection and laboratory processing

Following the discovery of a high percentage match between *Wolbachia* sequences and the BF3-BR2 primer pair (Elbrecht et al., 2019), three large amplicon sequencing datasets generated using these primers were investigated. The datasets, generated during the lab’s research on Swedish insect-associated microbiota, comprised amplicon data for both the COI and the bacterial 16S rRNA marker regions, and thus providing an independent source of information about *Wolbachia* presence.

The data consisted of wild-caught Swedish insect samples: 1) SPH (Swedish PHoridae) data from a recently published dataset of 1842 scuttle flies from six Malaise traps operated in June-July 2018 in Northern Sweden (Nowak et al., 2025); 2) SIP (Swedish Insects - Phoridae) data, comprising 927 individuals, representing diverse scuttle flies from Malaise traps operated during 2019 across Sweden, as a part of Insect Biome Atlas initiative (Miraldo et al., 2025); 3) SIN (Swedish INsects) data of taxonomically diverse 5132 insects other than phorids, originating from the same traps as the SIP collection (Supplementary Table S3).

Specifically, insects captured in fourteen (SIN, SIP - eight, SPH - six) different sites across Sweden, originally killed and stored in 95% ethanol (concentrations could have decreased during collecting, but were brought back to at least 80% once in the lab), were transported to Station Linné entomological station, and stored at -20 °C until processing. Then, trap contents were manually sorted into bins representing taxonomic groups representing Station Linné’s first tier (Forshage & Karlsson, n.d.; Karlsson et al., 2020), generally corresponding to orders and suborders, with an additional effort made to separate Phoridae from other Diptera. From these bins, we have selected morphologically distinct individuals to maximise the fraction of natural diversity processed. The collection dates and locations are included in the Supplementary Table S3.

Both SIN and SIP samples were processed using laboratory and bioinformatics pipelines detailed recently (Buczek et al., 2023; Nowak et al., 2025). Briefly, insects were individually homogenised using a lysis buffer (0.4 M NaCl, 10 mM Tris–HCl, 2 mM EDTA, 2% SDS, 10 µl proteinase K) and different-sized ceramic beads on a Bead Ruptor Elite homogeniser (Omni International). Then, 40 µl aliquots of samples were combined with a known number of plasmid copies carrying artificial 16S rRNA sequences (Tourlousse et al., 2016) for quantification purposes, and then purified on a 96-well plate using SPRI beads. The amplicon libraries were prepared following a 2-step PCR protocol: for the first step, we used a mixture of template-specific primers with Illumina adapter stubs, with the resulting PCR products used as a template for the second indexing PCR. Template-specific primers in PCR1 comprised a mixture including BF3-BR2 for insect COI (Elbrecht et al., 2019), 515F-806R for 16S-V4 rRNA (Parada et al., 2016), and in most cases, also primers for V1-V2 regions of 16S rRNA (Walker et al., 2015) or fungal ITS regions (Martin & Rygiewicz, 2005), not analysed in this study. The final libraries were verified on an agarose gel and pooled approximately equimolarly based on band intensities. The purified pool was combined with pools representing other projects and submitted to the Swedish National Genomics Infrastructure for sequencing on Illumina NovaSeq 6000 SPrime flow cells (2×250 bp read mode). To control for contamination from various sources, for each of the 88 biological samples processed, we used eight different controls, including negative controls for the DNA extraction, first PCR, and indexing PCR, as well as positive controls.

### Bioinformatic processing of the amplicon datasets

The raw sequencing reads for 8,798 samples were split into bins representing the COI and 16S-V4 rRNA targets based on primer sequences. The forward and reverse reads for each bin were merged, filtered based on length and quality, and then dereplicated, with singletons removed. Then, data for individual libraries were denoised, and the tables with information on the distribution of zero-radius Operational Taxonomic Units (zOTUs, a practical equivalent of Amplicon Sequence Variants - ASVs) across samples were reconstructed. Afterwards, zOTUs were clustered into 97% similarity operational taxonomic units (OTUs). Taxonomy assignment was done using the SINTAX algorithm and taxonomic reference databases: for the COI data, MIDORI version GB 239, with alphaproteobacterial sequences added; and for 16S rRNA data, SILVA version 138 SSU. For COI, sequences below 100% classification confidence score at the level of order (for insects) or domain (for *Wolbachia*) have been reclassified using the full NCBI nt database using the BLAST algorithm. For insect barcode assignment, we followed the workflow described previously (Nowak et al., 2025), but with altered parameters: specifically, we have assigned the samples’ barcode as the most abundant sequence whose taxonomy was classified as insect. Then, samples from SIN and SIP datasets were filtered out based on the following criteria: lack of 16S-V4 data (9 samples), lack of insect-assigned COI barcode (53 samples), fewer than 50 reads of barcode OTU (217), less than 90% dominance of the barcode OTU in sample (554 samples), lack of extraction spike-in reads (64 samples). This filtering yielded the final dataset of 7901 samples. The sequences representing *Wolbachia* were extracted from the full dataset based on their taxonomy assignment. Using these extracted reads, we compared the congruence of the *Wolbachia* signal between the two markers. The correlation plot of COI and 16S-V4 rRNA data for the samples with both markers’ signals has been generated in RStudio using the ggplot2 package v4.0.2 (Wickham, 2016).

### Comparison of *Wolbachia* detection reliability between COI and 16S-V4 rRNA

The overlap in *Wolbachia* detection between the two markers was evaluated across a range of thresholds. The parameters tested included the absolute number of *Wolbachia* reads per sample and the proportion of *Wolbachia* reads relative to the total reads within a given marker. For COI data, an additional parameter was tested, defined as the ratio of *Wolbachia* reads to insect barcode reads. To account for within-sample sequence diversity, each parameter was calculated in two ways: using either the read count of the most abundant *Wolbachia* sequence in a sample or the total number of *Wolbachia* reads across all sequences in that sample. For each of these parameters, a pairwise comparison between the marker genes was conducted using a dedicated script that scanned the thresholds providing the highest overlap in *Wolbachia* detection. The overlap was measured as a ratio of the number of samples assigned as *Wolbachia*-positive based on both markers, divided by the number of all samples with *Wolbachia* signal for at least one marker. To account for the taxonomic differences, the optimal parameters and threshold values were calculated separately for SPH dataset, SIP dataset, and insect orders within SIN dataset, excluding orders with fewer than 28 samples. The universal optimal threshold represents cutoff values maximising the mean overlap percentage among the abovementioned sample groups. Each group was weighted equally, and the universal values did not change when calculated for the SIN dataset alone.

### The correlation between COI and metagenomic *Wolbachia* signal and assessment of phylogenetic resolution provided by alternative datasets

Based on the presence and diversity of *Wolbachia* signal in 16S-V4 rRNA and COI data, we selected 152 diverse insect samples from the SIN dataset for metagenomic analysis. The samples were selected to cover diverse combinations of host and *Wolbachia* genotypes. Metagenomic libraries were prepared in-house using NEBNext Ultra II FS DNA Library Prep Kit, and the pools were submitted to Novogene Ltd. for sequencing using Illumina NovaSeq X Plus platform with 300-cycle flow cells, to an average coverage of 42 million reads per read pair. The raw reads were submitted to GenBank (PRJNA1470654), and the individual libraries are listed in the Supplementary Table S4.

To evaluate the correlation between *Wolbachia* signal in amplicon and metagenomic libraries, raw paired-end metagenomic reads were quality trimmed with automatic adapter detection and removal of poly-G tails of at least 5 bp. Then, these filtered reads were subsequently mapped against sample-specific host barcodes and dominant *Wolbachia* amplicon sequences, using minimap2 v2.24-r1122 (Li, 2018) in short-read mode (-ax sr). Alignments were filtered with SAMtools v1.9 (Danecek et al., 2021) to retain reads with mapping quality of at least 20 while excluding unmapped, secondary, and supplementary alignments. Afterwards, the *Wolbachia*-to-host coverage ratio was calculated by dividing the mean sequencing depth of the *Wolbachia* COI reference by the depth of the host COI barcode sequence, as estimated using samtools depth. After a log10 transformation, the correlation between metagenome- and amplicon-based *Wolbachia*-to-host coverage ratios was checked using Spearman’s test and visualised in RStudio using the ggplot2 v4.0.2 package (Wickham, 2016).

To assess phylogenetic resolution provided by the COI amplicon reads, conserved-length *Wolbachia* protein-coding genes were identified from the previously mentioned Prokka annotation of reference genomes and extracted from metagenomes. To do so, raw reads were quality filtered with a Phred quality threshold of 30, a maximum error rate of 0.15, and a minimum retained read length of 80 bp. The filtered reads were subsequently processed using HybPiper 2.3.0 (Johnson et al., 2016) in DNA mode, using a custom *Wolbachia* reference dataset comprising phylogenetically informative genes mentioned before. Read mapping and target recovery were conducted with the BWA implemented in HybPiper, using a minimum coverage cutoff of 8× for sequence assembly. The recovered DNA sequences were extracted with HybPiper retrieve_sequences to obtain stitched contigs. The gene was assigned as ‘recovered’ if its stitched contigs’ length was >80% of the reference gene. After a manual assessment of the recovery success, libraries containing fewer than 80 out of 153 genes were discarded. For the 87 samples with at least 80 of

153 genes successfully recovered, genes were concatenated and then used to construct a phylogenetic tree using IQTREE 3.0.1 (Wong et al., 2026). The phylogeny was visualised with iTOL v7 (Letunic & Bork, 2007). For the same set of samples, amplicon sequences of dominant *Wolbachia* genotypes 16S-V4 rRNA and COI markers were visualised using TCS (Snell et al., 2002) haplotype networks from PopArt 1.7 (Leigh & Bryant, 2015) and their mean pairwise genetic distance calculated using MEGA version 11.0.13 with the p-distance model (Tamura et al., 2021).

## Results

### Standard COI barcoding primers match *Wolbachia* and diverse other bacteria

We examined the extent to which *Wolbachia* cytochrome oxidase I sequences match commonly used insect barcoding primers targeting different portions of the Folmer region of this gene (Fig. 1). Across this region, we compared 56 established primers (28 forward and 28 reverse), together with the commonly used forward primer for *Wolbachia* multilocus sequence typing (Baldo et al., 2006). These primers were evaluated against 73 unique *Wolbachia* COI sequences representing thirteen currently available supergroups (Supplementary Fig. S1, Supplementary File S1).

**Fig. 1.**
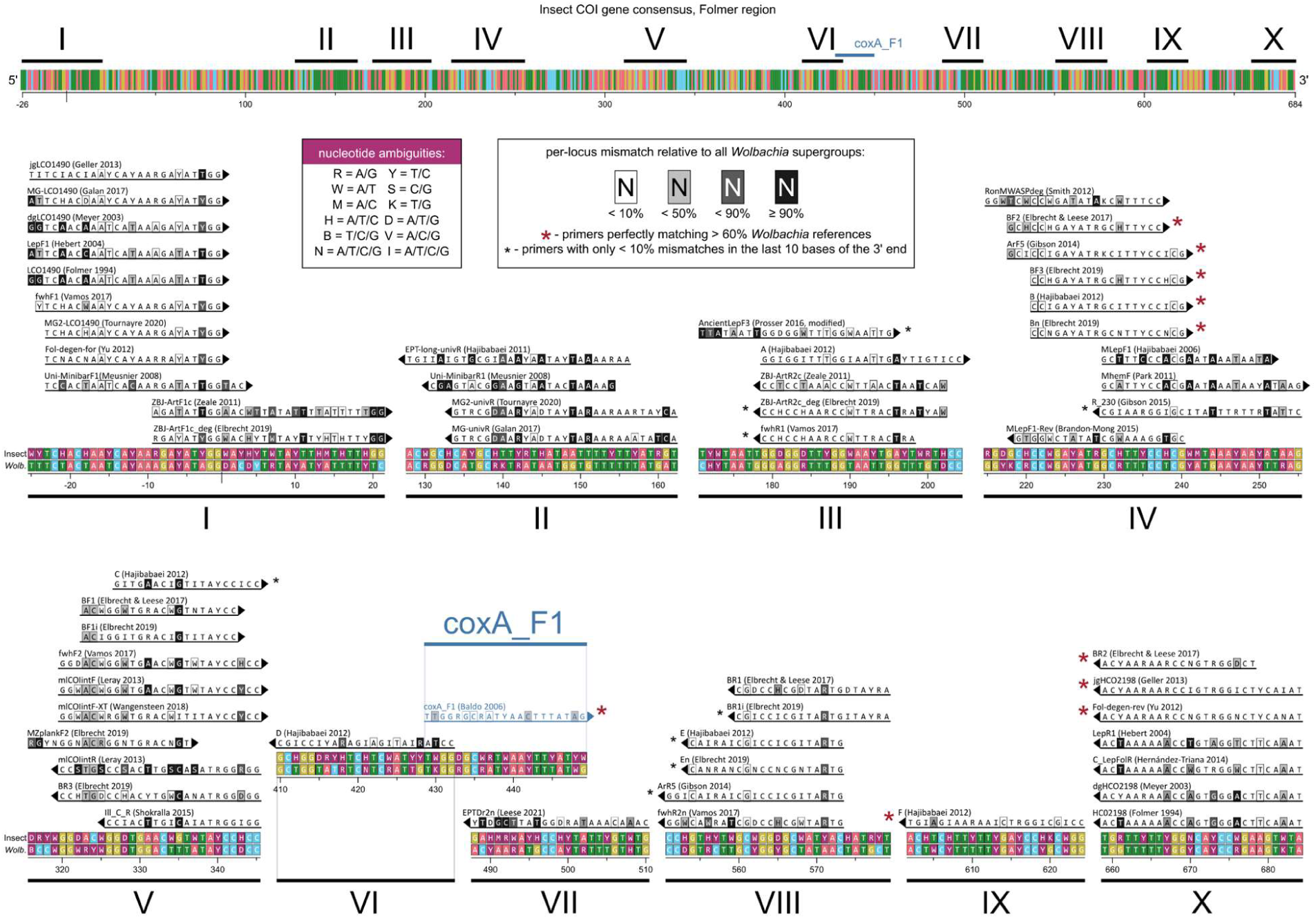
*In silico* evaluation of commonly used insect COI primers for potential amplification of *Wolbachia* sequences. The alignment of 56 invertebrate COI barcoding primers and the canonical *Wolbachia* MLST gene coxA forward primer (coxA_F1) within ten largely conserved portions (I-X) of the insect COI Folmer region and the homologous *Wolbachia* region. For each nucleotide position within each primer sequence, information is provided on the percentage of mismatches relative to all *Wolbachia* reference sequences.

Among the insect primers tested, five forward and four reverse primers showed perfect matches, when allowing for degenerate positions, to the majority (>50%) of the reference *Wolbachia* sequences (Supplementary Table S5). Two reverse primers matched all available *Wolbachia* sequences without any mismatches: Fol-degen-rev (Yu et al., 2012) and jgHCO2198 (Geller et al., 2013). Interestingly, forward primer B (Hajibabaei et al., 2012) perfectly matched more reference *Wolbachia* sequences than the dedicated coxA_F1 primer for *Wolbachia* multilocus sequence typing, which had mismatches for six out of thirteen tested supergroups. Two further forward primers and seven reverse primers had <10% mismatch to reference *Wolbachia* sequences within the first ten nucleotides at the 5′ end of the primer, most critical for primer annealing and the initiation of extension, and were thus likely to support amplification at least partially. In contrast, many other primers exhibited substantial mismatches, frequently located within the first few nucleotides at the 5′ end.

Based on this analysis, popular insect barcoding primer pairs such as BF3–BR2 are predicted to amplify more than 90% of *Wolbachia* sequences examined. In contrast, barcoding assays relying on primers such as LCO1490 and HCO2198, mlCOIintF and mlCOIintR, or fwhF2 and fwhR1, as well as their derivatives, are expected to yield considerably lower amplification success for *Wolbachia* sequences. The results, however, varied among *Wolbachia* supergroups. Nine insect barcoding primers matched most sequences belonging to supergroups A and B. At the same time, fourteen primers showed perfect matches to at least one sequence from supergroup D and only six primers matched sequences from supergroup F. These results indicate substantial variation in primer compatibility across the phylogenetic diversity of *Wolbachia* (Supplementary Table S5).

Diverse bacterial sequences other than *Wolbachia* can also be matched by insect COI primers (Table 1, Supplementary Table S6). We found that 21,824 of 59,318 (36.8%) of bacterial RefSeq genomes, particularly in phyla Pseudomonadota and Actinomycetota, encode at least one copy of *ctaD* gene – the COI homolog. 25.0% of these genomes encoded more, and up to eight, distinct *ctaD* copies. A surprisingly large share of these genes was matched by some insect barcoding primers, with 24 of the 57 tested primers matching at least some bacterial references (Table 1, Supplementary Table S6). For example, jgHCO2198 perfectly matched *ctaD* copies in 17,410 (79.8%) of genomes that encoded it, and a partly overlapping BR2 matched 15,397 (70.6%). These reverse primers, located at around positions 660-680 of insect COI gene, had broad taxonomic coverage spanning Pseudomonadota and Actinomycetota. The forward primers were more selective and generally restricted to a subset of Alphaproteobacteria. For example, BF3 perfectly matched only 413 RefSeq genomes (1.9%), the large majority of which were Rickettsiales (incl. *Wolbachia*). Interestingly, when multiple distinct *ctaD* variants were present in a genome, they often matched the same COI primers.

**Table 1.**
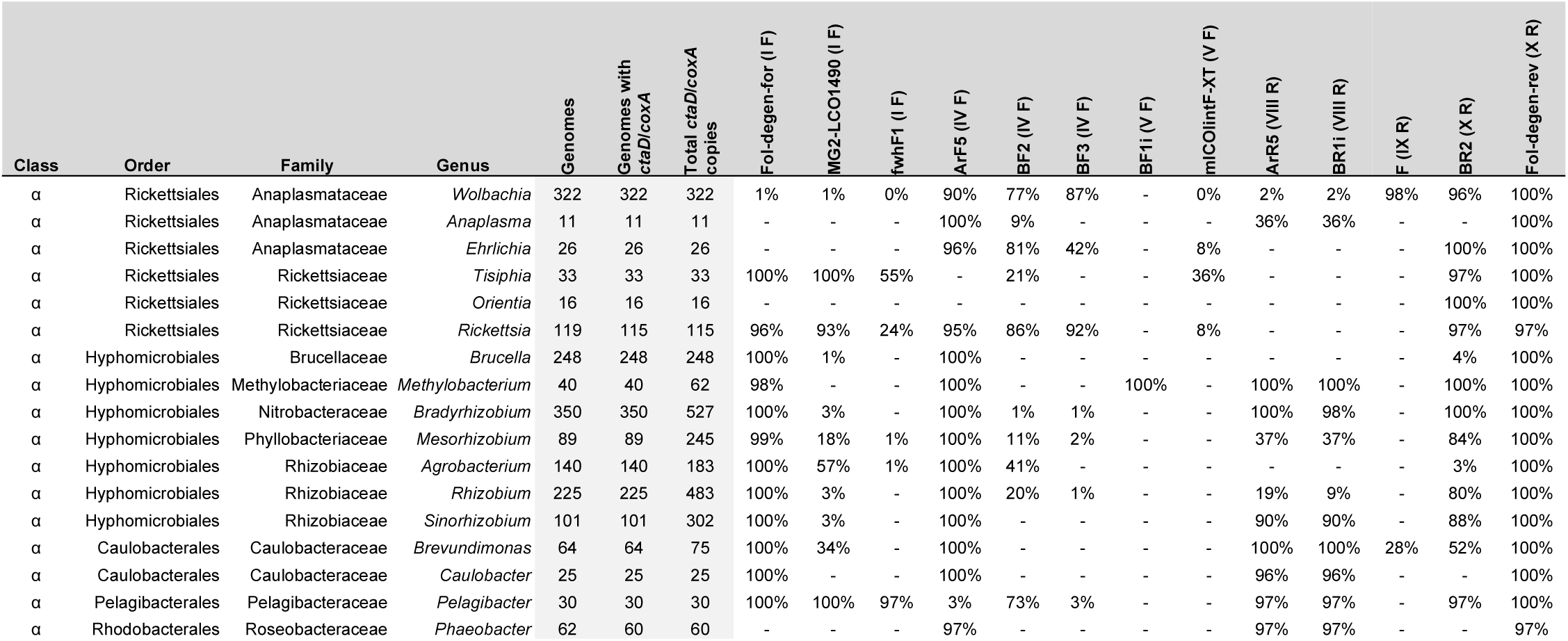
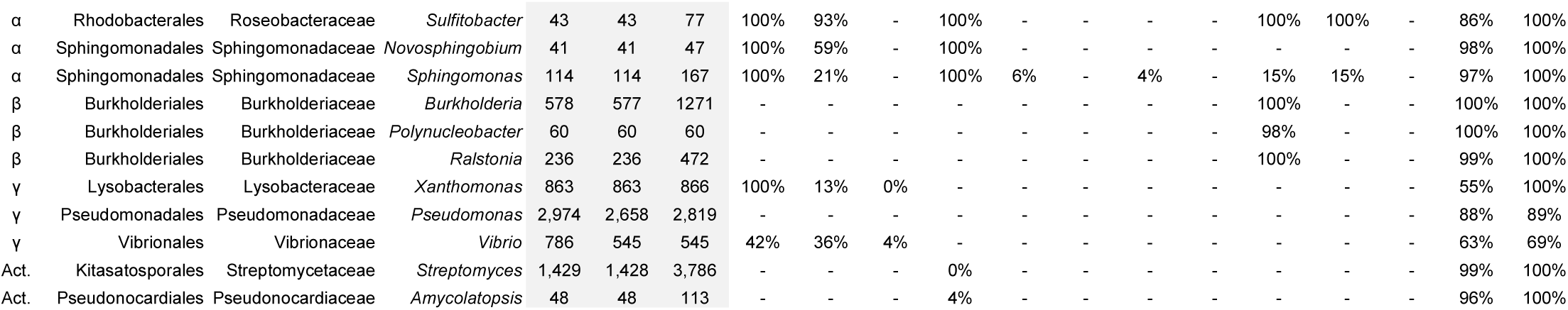
Selected bacterial genera well-represented in the NCBI RefSeq collection, with information on *coxA/ctaD* gene copies they carry, and the percentage of genomes with at least one gene copy perfectly matching selected COI insect barcoding primers. Abbreviations for bacterial classes: α - Alphaproteobacteria, β - Betaproteobacteria, γ - Gammaproteobacteria, Act. - Actinomycetes. For primers, the number in brackets indicates the COI gene region, as indicated in Figure 1, and F/R - their direction.

### Wolbachia is often present in COI amplicon data

To evaluate the occurrence of *Wolbachia* sequences in insect barcoding data, we screened three multitarget amplicon datasets, representing 7,901 individual insects. For all of them, the COI region was amplified using popular BF3-BR2 primers, simultaneously with the V4 region of the 16S rRNA gene. This included 1,842 Northern Swedish scuttle flies (recently published SPH dataset (Nowak et al., 2025)), 927 further Swedish scuttle flies (SIP dataset), and a taxonomically diverse dataset comprising 5,132 various insects (SIN), as detailed in the Methods. To accommodate the taxonomic breadth of the material, results for the last dataset were analysed separately for insect orders (Supplementary Table S7, Supplementary File S2).

*Wolbachia* sequences were common within COI data across all three datasets. The COI libraries yielded a median of 20,654 total reads per sample (average 28,418, IQR: 7,217-41,329). Most reads corresponded to insect barcode, contributing a median of 19,685 reads (average 27,396, IQR: 6,727-40,072) or 98.7% of the total reads per sample (average 94.6%, IQR: 94.9%-99.9%). After singleton removal, *Wolbachia* COI reads were found in 35.1% of the samples (Fig. 2a). When present in a sample, *Wolbachia* contributed a median of 221 COI reads (average 708, IQR: 49-731), corresponding to a median of 1.8% of reads (average 6.7%, IQR: 0.3%-6.5%) per COI library. The largest share of *Wolbachia* reads was found among Hymenoptera, where in the 29.4% of samples where their reads were present, *Wolbachia* accounted for a median of 14.3% of the total reads (average 23.7%, IQR: 4.3%-36.8%) (Fig. 2b). This was expected, as some Hymenoptera are known to amplify less efficiently with commonly used COI barcoding primers because of primer–template mismatches (Elbrecht et al., 2019). The frequency of samples containing *Wolbachia* COI reads varied substantially among other insect taxa as well. In both scuttle fly datasets, the fraction exceeded 45%, whereas in several less represented insect orders, including the few processed Plecoptera and Trichoptera, no *Wolbachia* COI reads were recovered (Fig. 2a). Among the *Wolbachia*-positive samples, in 32.0% of cases, more than one COI *Wolbachia* sequence variant was found per library (Supplementary Table S7). This likely represents a mixture of biological signal and methodological artifacts - a combination of superinfections with pseudogenes, cross-contamination among samples through index hopping, and sequencing noise. Nevertheless, when reads of multiple *Wolbachia* zOTUs were present in a library, one zOTU almost always comprised their majority (average 77.9%, median 82.2%, IQR: 62.3-96.6%), in most cases representing the true symbiont, with lower confidence regarding the additional variants. For simplicity, further analyses rely on the most abundant *Wolbachia* sequence per sample. These most abundant sequences accounted for a median of 1.7% (average 5.9%, IQR: 0.3%-5.6%) of the total reads per *Wolbachia*-positive sample.

**Fig. 2.**
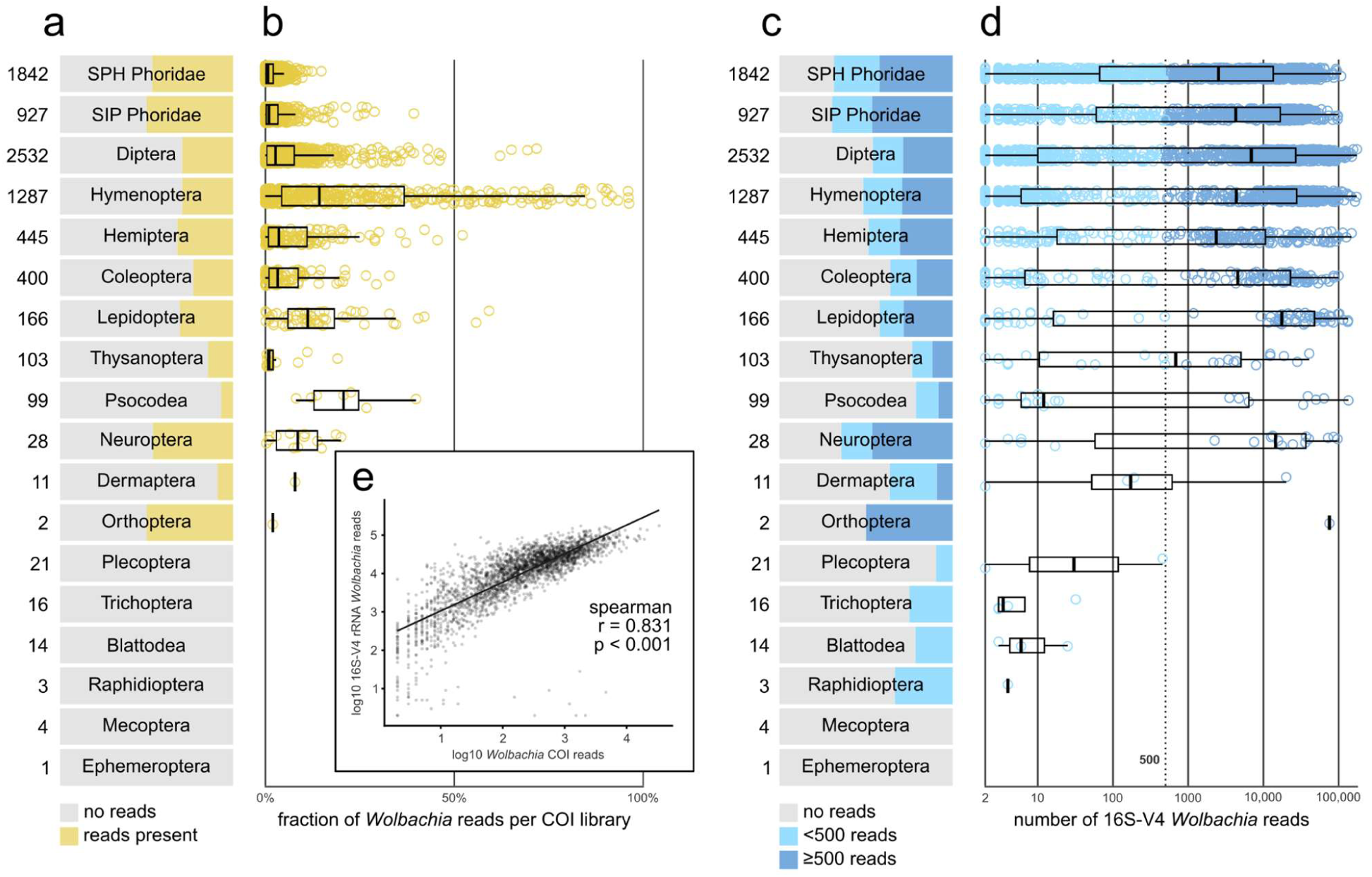
The prevalence and relative abundance of amplicon reads corresponding to *Wolbachia* in COI amplicon and 16S-V4 rRNA amplicon data across 7,901 taxonomically diverse individual insects. a) The proportion of samples with at least 2 *Wolbachia* COI reads across insect taxonomic bins. b) Relative abundance of *Wolbachia* reads in COI libraries across the datasets. c) The proportion of samples with at least 2 *Wolbachia* reads in 16S-V4 rRNA data across insect taxonomic bins. d) Number of per-sample *Wolbachia* 16S-V4 rRNA reads across the datasets. e) The correlation among the numbers of *Wolbachia* COI and 16S-V4 rRNA reads in two-gene amplicon datasets for the same samples. The cutoff of five hundred reads for the 16S-V4 rRNA data was selected as an arbitrary threshold above which we considered *Wolbachia* presence reliable.

For the simultaneously amplified 16S-V4 rRNA, about 53.8% of samples contained at least two *Wolbachia* reads, which was a noticeably higher percentage than 35.1% in the COI data (Fig. 2c). 16S-V4 rRNA marker data for these same 7,901 insects (Supplementary Table S7, Supplementary File S3) contained a median of 14,548 reads (average 27,732, IQR: 3,611-40,356), after discarding synthetic quantification spike-in reads that were not used in analyses. In many of these cases, the 16S-V4 rRNA read counts were low (median 8, average 850, IQR: 4-30) (Fig. 2d), and similarly to COI dataset, they may often represent different sources of background noise rather than true symbioses; in particular, cross-contamination through index hopping is likely to be more pronounced when a large share of samples sequenced in the same flow cell harbour high-abundance *Wolbachia* (Van Der Valk et al., 2020). Despite a discrepancy in detection frequency between the two markers, *Wolbachia* read abundance showed a significant positive relationship among these markers, with higher COI read counts associated with higher 16S rRNA counts (Fig. 2e).

Applying an arbitrary threshold of five hundred reads in the 16S rRNA dataset, unlikely to be achieved through cross-contamination and other artifacts alone, substantially increased agreement between the two markers, producing a strong congruence between *Wolbachia* detection based on COI and 16S rRNA data.

### *Wolbachia* is reliably detected using COI amplicons when 16S-V4 rRNA data is used as a reference

We evaluated whether *Wolbachia* presence inferred from COI barcoding data is consistent with detection based on the universal bacterial 16S-V4 rRNA marker. The overlap in *Wolbachia* presence between the two markers, with different abundance thresholds, was quantified across samples. All zOTUs present in a library only once had been discarded at the data filtering stage; and when the occurrence of any *Wolbachia* zOTU represented by >1 read was treated as evidence of presence, the agreement between markers was limited. Under this permissive definition, in 54.5% of samples, *Wolbachia* reads were present for at least one marker. However, in 37.0% of these cases, *Wolbachia* was represented in one marker only (Fig. 3a). Much of the discrepancy involved samples with low *Wolbachia* read counts within the 16S-V4 rRNA dataset and none in the COI dataset. To reduce the influence of low-level signals that we considered less reliable, we applied more stringent thresholds. For the 16S-V4 rRNA data, we used the universal cutoff of >324 reads of the most abundant *Wolbachia* zOTU. For the COI data, with a much more variable total number of reads, we used relative abundance threshold, where the ratio of dominant *Wolbachia* sequence reads to insect barcode reads was greater than 0.000194. These thresholds were selected empirically by maximising marker agreement across the datasets and insect orders. Under these criteria, the congruence between the two markers increased substantially, exceeding 90% in most insect groups (Fig. 3b) and in many cases approaching the theoretical maximum achievable for the datasets (Fig. 3c). Nevertheless, the optimal thresholds varied among taxa. For example, the best 16S rRNA threshold for Coleoptera exceeded 1,000 reads, whereas in the SPH scuttle fly dataset, a higher COI proportion threshold than the universal value was required to achieve the maximal agreement (Fig. 3d). Across all taxa, however, the taxon-optimal thresholds did not increase the overlap by more than 3.1 percentage point relative to the universal threshold.

**Fig. 3.**
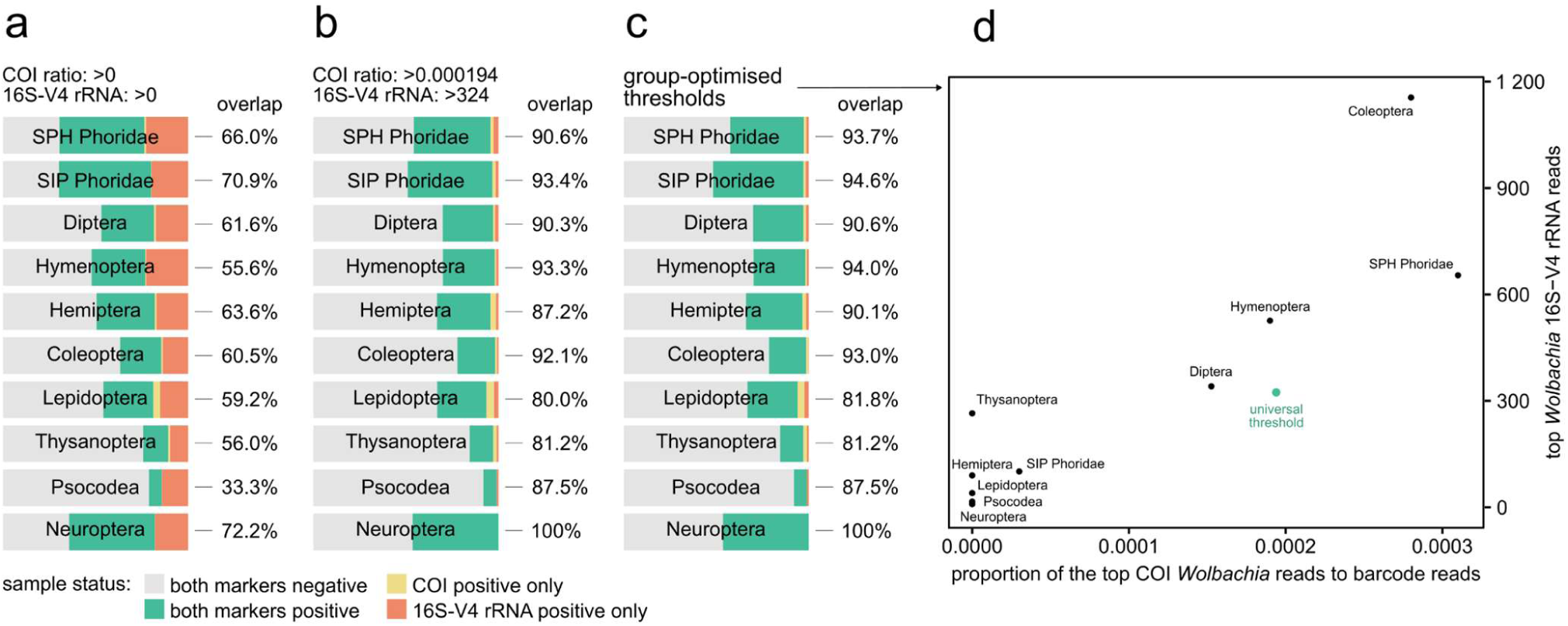
Overlap between COI and 16S-V4-based *Wolbachia* presence inference under alternative data filtering strategies. Detection based on 16S-V4 rRNA data was inferred using the read count of the most abundant *Wolbachia* zOTU per sample. Detection based on COI data was inferred using the ratio of reads assigned to the dominant *Wolbachia* zOTU relative to reads assigned to the host barcode zOTU. Overlaps are shown for three filtering schemes: a) no filtering, b) universal cut-offs optimised across all datasets, and c) dataset-specific cut-offs. d) Distribution of the optimal cut-off values for datasets and insect orders.

### *Wolbachia* COI amplicon data correlate with the corresponding signal in metagenomic libraries

Given the strong correspondence in *Wolbachia* detection between the COI and 16S-V4 amplicon datasets, we next evaluated whether the abundance of *Wolbachia* reads in amplicon libraries reflects the strength of the signal in shotgun metagenomic data. To this end, metagenomic reads from 152 insect specimens were mapped against the amplicon-reconstructed partial COI sequences of the host and of the dominant *Wolbachia* in the respective samples. After comparing the relative coverages of said sequences in each sample, we observed a strong positive correlation between the *Wolbachia*-to-host ratios among amplicon and metagenomic datasets for the same insects (Fig. 4a). Samples with high *Wolbachia*-to-host amplicon read ratio (>0.02) most often produced high metagenomic coverage (median 27×, average 47×, IQR: 13-44×), whereas specimens with low amplicon-derived signal (ratio <0.0005) generally presented coverage below 1× or did not map at all in the corresponding metagenomic libraries (Supplementary Table S8). These results indicate that *Wolbachia* abundance inferred from standard amplicon datasets might broadly reflect underlying biological variation rather than amplification biases.

**Fig. 4.**
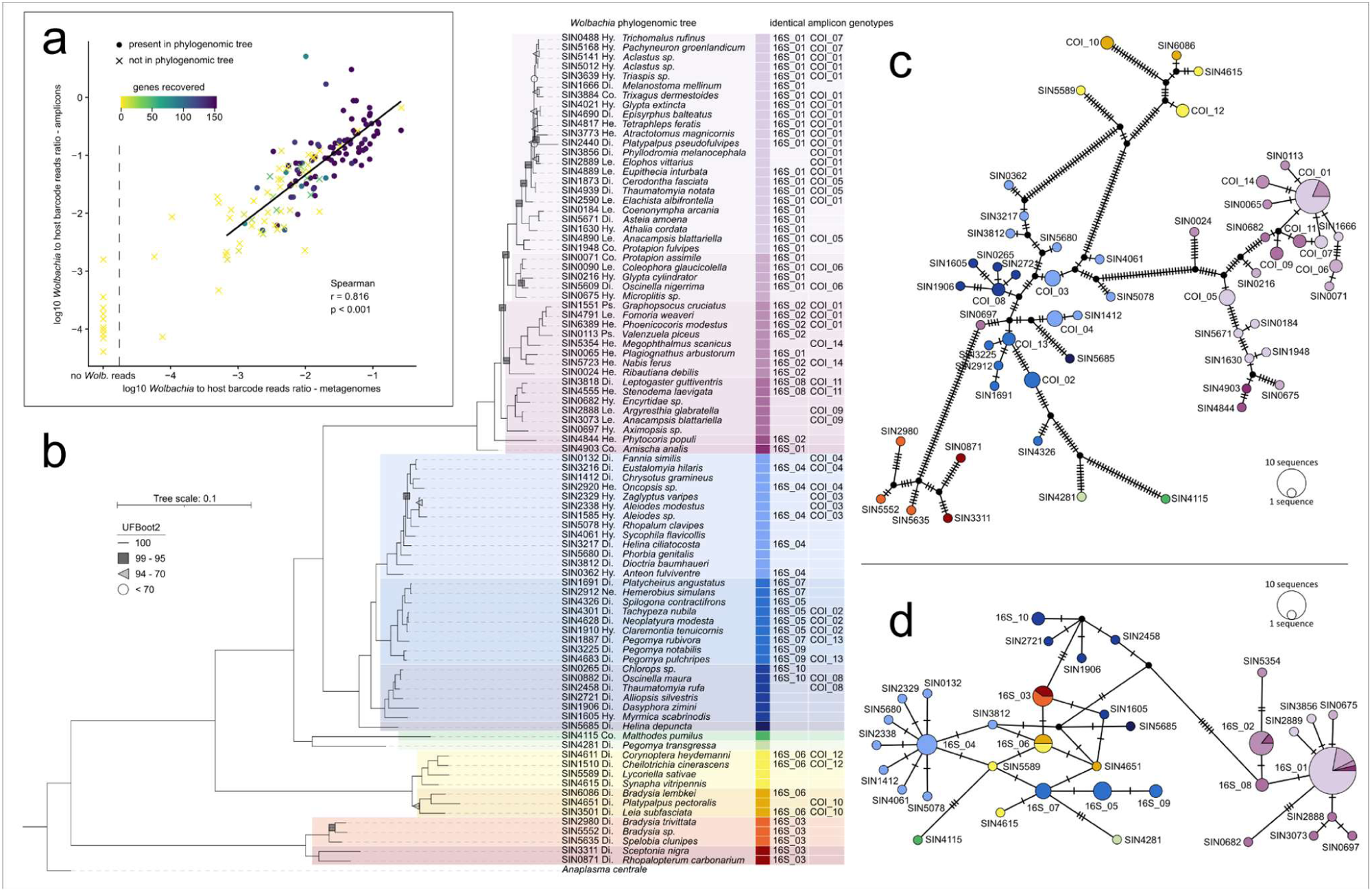
Comparison between the genomic tree and amplicon read-based *Wolbachia* networks a) Correlation between *Wolbachia*-to-host read number ratios among COI amplicon and metagenomic datasets for 152 individual insects; b-d) Phylogenetic relationships inferred for: b) 87 samples based on a 153-gene dataset (121 kbp), c) 55 COI haplotypes of 418 bp, and d) 36 16S-V4 rRNA haplotypes of 253 bp, that these 87 samples represented.

Apart from the amplicon mapping approach, we have also used the metagenomic data to extract 153 phylogenetically informative genes for further analysis. The success of marker gene recovery from metagenomic data was again positively associated with *Wolbachia*-to-host read number ratios in amplicon data. Notably, samples where at least eighty genes had the completeness of >80%, and that were used for the phylogenomic tree reconstruction, had the highest *Wolbachia*-to-host ratio among amplicon libraries (average 0.27, mean 0.11, IQR: 0.05, 0.30). Samples for which we recovered some but fewer than eighty phylogenetically informative genes had significantly lower ratios (average 0.02, median 0.03, IQR: 0.01, 0.05) and samples which did not yield usable genes often had the lowest amplicon ratio (average 0.01, median 0.06, IQR: 0.002, 0.04) (Fig. 4a, Supplementary Table S8).

### Wolbachia COI amplicon data provide meaningful phylogenetic resolution

Using metagenomic data, we have generated a genome-based phylogeny for strains from 87 *Wolbachia*-positive insect specimens where we succeeded in reconstructing at least eighty phylogenetically informative genes. The genome-based phylogeny revealed at least five deeply divergent clades (Fig. 4b: purple, blue, green, yellow, red), comprising multiple genotypes each. This phylogeny was compared with a genotype network constructed based on amplicon sequences alone. For COI, despite clustering of closely related clades, the overall concordance with the genomic tree remained high. The 87 COI sequences included in the comparison represented 55 unique haplotypes and displayed substantial divergence (mean pairwise genetic distance 9.6%), clearly separating all five major clades (Fig. 4c). Based on the previously generated phylogeny using the 709 bp Folmer region, which almost perfectly separated *Wolbachia* supergroups, these clades are likely to represent supergroup level (Supplementary Fig. S1). In contrast, the sequences for the more conserved 16S-V4 rRNA marker represented 36 genotypes and had a considerably lower resolution (mean pairwise genetic distance of 2.8%). While it separated the supergroup B and distinguished some strains, the extensive sequence similarity among genotypes obscured much of the underlying phylogenetic structure (Fig. 4d).

## Discussion

Our study demonstrates that standard insect COI barcoding datasets frequently contain biologically informative *Wolbachia* sequences. Because several widely used invertebrate COI primer pairs closely match homologous *coxA*/*ctaD* regions of *Wolbachia* and other bacteria, co-amplification is a widespread feature of contemporary barcoding workflows. With proper planning, this creates promising opportunities to investigate host–symbiont associations across large insect collections.

*Wolbachia* recovery from COI data is highly primer dependent. While many commonly used COI primers showed substantial mismatches against *Wolbachia* reference sequences, several widely adopted primer pairs, including BF3-BR2, matched *Wolbachia* references with few or no mismatches, sometimes surpassing a dedicated *Wolbachia*-targeting primer. Consequently, retrospective mining of existing barcoding datasets for *Wolbachia* signal should prioritise datasets generated using the most compatible primers. Nevertheless, *in silico* primer analysis alone cannot fully predict amplification performance under laboratory conditions (Elbrecht et al., 2019; Elbrecht & Leese, 2017). The efficiency of *Wolbachia* co-amplification is also expected to depend on PCR conditions, including primer annealing temperature and polymerase chemistry, as well as on the relative abundance of host and symbiont DNA (Elbrecht et al., 2019; Piñol et al., 2015, 2019). Interestingly, while primers also match many other bacteria known to colonise insects, we have not often encountered their reads in COI amplicon data. This may be because of their lower prevalence and abundance, or our team’s long-term commitment to the use of Rickettsiales-restricted primer BF3; hence, it would be valuable to inspect data obtained with alternative primers.

Although our estimates are based on individual specimens rather than species, the overall prevalence is comparable to previous estimates of *Wolbachia* incidence across insects (Kaur et al., 2021; Weinert et al., 2015). However, because primer performance varied among *Wolbachia* clades, some lineages are likely to be preferentially amplified while others remain underrepresented. This mirrors the well-recognised limitation of insect barcoding itself, where “universal” primers differ in amplification efficiency across taxonomic groups (Elbrecht & Leese, 2017). Therefore, comparisons of *Wolbachia* prevalence or abundance among datasets generated using different protocols should be interpreted cautiously, given the variation in primer performance in both insects and *Wolbachia*. At the same time, while amplification efficiency is expected to vary among markers, taxa, and laboratory protocols, the strong overlap between COI- and 16S rRNA-based *Wolbachia* detection in our study suggested that this did not obscure major patterns in our datasets.

Critically, however, the detection of *Wolbachia*-derived sequences, whether COI, 16S rRNA, or others, should not automatically be interpreted as evidence of stable heritable association with the focal insect host. Cases where we see the same abundant *Wolbachia* genotypes in a significant subset of individuals within a population often indicate a true established symbiotic association. Other cases are less clear. For example, some individuals may have been only recently colonised by a *Wolbachia* strain that spread across tissues but would not have been transmitted to offspring (Bright & Bulgheresi, 2010). Interpretation of weak signals may be particularly challenging, as apart from real, low-titre presence, *Wolbachia* DNA, or reads, may originate from parasites or parasitoids associated with the sampled insect, from recently consumed dietary material, from environmental or laboratory contamination, or from low-level index hopping during sequencing (Kolasa et al., 2023; Pompanon et al., 2012; Salter et al., 2014; Van Der Valk et al., 2020). In addition, fragments of the *Wolbachia* genome are known to integrate into host nuclear genomes in some arthropod lineages, potentially contributing to the signal (Song et al., 2008). These alternative scenarios are likely to explain some of the cases where few *Wolbachia* COI reads were seen. Consequently, COI-based *Wolbachia* detection is best viewed as a scalable screening approach that identifies candidate symbioses and broad ecological patterns, while definitive confirmation of stable symbioses would require complementary approaches such as microscopy, laboratory cultures, or dedicated genomic analyses. For the same set of reasons, we conclude that it would be very hard, if possible at all, to be absolutely certain about the distribution of heritable *Wolbachia* symbioses in a large collection of diverse wild-caught insects, regardless of the method used.

Beyond simple presence–absence inference, the recovered COI sequences provided valuable information regarding the phylogenetic placement of *Wolbachia* strains, providing considerably higher resolution than partial 16S rRNA sequences. *Wolbachia* COI genotype networks broadly reproduce relationships inferred from metagenomic data, generally allowing identification of major *Wolbachia* clades and discriminating among related genotypes. This outcome is consistent with the greater evolutionary rate and higher sequence variability of protein-coding genes relative to conserved ribosomal markers. At the same time, phylogenetic inference based on a single marker has well-recognised limitations, as recombination, horizontal gene transfer, and differences in evolutionary histories among loci may obscure relationships among *Wolbachia* strains (Baldo et al., 2006; Bleidorn & Gerth, 2018). Nevertheless, COI genotype resolution remains sufficiently informative to identify the main clades, detect previously unknown variants or polymorphisms within populations, and distinguish between widespread shared symbioses and putative independent acquisition events (Nowak et al., 2025). Specifically, identification of shared or highly similar *Wolbachia* haplotypes across distinct hosts suggests recent host shifts or circulation of related strains within ecological networks, whereas an observation of divergent haplotypes within a population or species would indicate independent symbiosis histories. At the same time, the discovery of the same or very similar symbiont genotypes across populations of a species, or closely related species, suggests stable associations, but would require confirmation using methods that provide higher phylogenetic resolution. Such population-level COI-distribution patterns may become particularly informative when interpreted together with ecological information on host identity, trophic interactions, parasitoid associations, or geographic distribution (Nowak et al., 2025). The ability to recover preliminary strain-level information directly from routine barcoding data for large numbers of diverse wild insects, and combine them with host information, creates opportunities to investigate these processes across spatial and taxonomic scales that would be difficult to achieve using other means.

More broadly, our findings illustrate that standard barcoding datasets may become an extensive source of information extending beyond species identification. Better understanding of the off-target amplification may improve barcoding pipelines and increase barcode reconstruction success, while reducing the risk of specimen misidentification. Alongside *Wolbachia*, insect COI datasets may also recover sequences from other bacteria possessing homologous oxidase genes, as well as from parasitoids, pathogens, dietary organisms, and other ecological associates (Hrček & Godfray, 2015; Kolasa et al., 2023; Pompanon et al., 2012). These secondary signals, rarely considered, may provide valuable information on ecological interactions and hidden dimensions of community structure. Developing approaches capable of systematically extracting and interpreting such information may considerably increase the research value of the ongoing, large-scale biodiversity monitoring programmes. Our results suggest that these rapidly expanding datasets can be mined to investigate symbiont diversity and host–microbe associations at unprecedented scales, without additional sequencing costs, increase in laboratory workload, or - depending on primers used - changes to existing protocols. Such approaches may facilitate analyses of *Wolbachia* symbiosis dynamics, identification of candidate host shifts, and reconstruction of broad transmission networks at ecosystem scales, alongside prioritisation of specimens for more detailed metagenomic investigation. Hence, integrating the careful inspection of off-target reads, for *Wolbachia* and other organisms, into barcoding-based biodiversity monitoring schemes could be a game-changer for our understanding of how host–microbe interactions contribute to the structure, function, and evolution of insect communities.

## Author contributions

KHN and PŁ conceived the ideas and designed methodology; MB, MM and MP-F processed samples and collected data; KHN and CV analysed the data; JD and DS contributed methodology; KHN and PŁ led the writing of the manuscript. All authors contributed critically to the drafts and gave final approval for publication.

## Acknowledgements

We are thankful to William R. Conner from the University of Montana for his substantial help with Prokka annotation pipeline. This project was supported by the Polish National Science Centre grants 2018/31/B/NZ8/01158 and 2021/43/B/NZ8/03376, by the Knut and Alice Wallenberg Foundation grant KAW2017.088, as well as a grant from the Priority Research Area BioS under the Strategic Programme Excellence Initiative at Jagiellonian University.

## Supplementary item legends

- Supplementary Figure S1: Full alignment of the analysed COI primers and *Wolbachia* references
- Supplementary Table S1: List of the accession numbers for *Wolbachia* reference sequences
- Supplementary Table S2: List of insect and *Wolbachia* COI primers tested
- Supplementary Table S3: Sample collection details and metadata of the newly generated amplicon and metagenomic libraries
- Supplementary Table S4: GenBank references of the newly generated metagenomic data
- Supplementary Table S5: Percentage of matches between selected COI primers and various *Wolbachia* supergroups
- Supplementary Table S6: Summary of the COI barcoding primers matches to bacteria other than *Wolbachia*
- Supplementary Table S7: Amplicon read counts in host and *Wolbachia* categories across all the samples
- Supplementary Table S8: Comparison of the *Wolbachia* signal between metagenomic and amplicon libraries
- Supplementary File S1: Insect and *Wolbachia* COI reference sequences and *Wolbachia*
supergroups consensus sequences
- Supplementary File S2: Read count table of sequences classified as *Wolbachia* in COI data
- Supplementary File S3: Read count table of sequences classified as *Wolbachia* in 16S-V4 rRNA data

## Data availability statement

Amplicon data available in the supplementary files. Raw metagenomic reads are available in GenBank at PRJNA1470654. All supplementary data and scripts are also available in the GitHub repository: https://github.com/KarolHub/Wolbachia-in-COI-data

## References

Abdala-Roberts, L., Puentes, A., Finke, D. L., Marquis, R. J., Montserrat, M., Poelman, E. H., Rasmann, S., Sentis, A., Van Dam, N. M., Wimp, G., Mooney, K., & Björkman, C. (2019). Tri-trophic interactions: Bridging species, communities and ecosystems. Ecology Letters, 22(12), 2151–2167. 10.1111/ele.13392

Baldo, L., Dunning Hotopp, J. C., Jolley, K. A., Bordenstein, S. R., Biber, S. A., Choudhury, R. R., Hayashi, C., Maiden, M. C. J., Tettelin, H., & Werren, J. H. (2006). Multilocus sequence typing system for the endosymbiont *Wolbachia pipientis*. Applied and Environmental Microbiology, 72(11), 7098–7110. 10.1128/AEM.00731-06

Bleidorn, C., & Gerth, M. (2018). A critical re-evaluation of multilocus sequence typing (MLST) efforts in *Wolbachia*. FEMS Microbiology Ecology, 94(1). 10.1093/femsec/fix163

Bleidorn, C., & Henze, K. (2021). A new primer pair for barcoding of bees (Hymenoptera: Anthophila) without amplifying the orthologous coxA gene of *Wolbachia* bacteria. BMC Research Notes, 14(1), 427. 10.1186/s13104-021-05845-9

Bright, M., & Bulgheresi, S. (2010). A complex journey: Transmission of microbial symbionts. Nature Reviews Microbiology, 8(3), 218–230. 10.1038/nrmicro2262

Buchner, D., Sinclair, J. S., Ayasse, M., Beermann, A. J., Buse, J., Dziock, F., Enss, J., Frenzel, M., Hörren, T., Li, Y., Monaghan, M. T., Morkel, C., Müller, J., Pauls, S. U., Richter, R., Scharnweber, T., Sorg, M., Stoll, S., Twietmeyer, S., … Leese, F. (2025). Upscaling biodiversity monitoring: Metabarcoding estimates 31,846 insect species from Malaise traps across Germany. Molecular Ecology Resources, 25(1), e14023. 10.1111/1755-0998.14023

Buczek, M., Prus-Frankowska, M., & Łukasik, P. (2023). Quantitative multi-target amplicon sequencing workflow v1. 10.17504/protocols.io.36wgq351ylk5/v1

Chua, P. Y. S., Bourlat, S. J., Ferguson, C., Korlevic, P., Zhao, L., Ekrem, T., Meier, R., & Lawniczak, M. K. N. (2023). Future of DNA-based insect monitoring. Trends in Genetics, 39(7), 531–544. 10.1016/j.tig.2023.02.012

Cogni, R., Ding, S. D., Pimentel, A. C., Day, J. P., & Jiggins, F. M. (2021). *Wolbachia* reduces virus infection in a natural population of *Drosophila*. Communications Biology, 4(1), 1327. 10.1038/s42003-021-02838-z

Cruaud, P., Rasplus, J.-Y., Rodriguez, L. J., & Cruaud, A. (2017). High-throughput sequencing of multiple amplicons for barcoding and integrative taxonomy. Scientific Reports, 7(1), 41948. 10.1038/srep41948

Danecek, P., Bonfield, J. K., Liddle, J., Marshall, J., Ohan, V., Pollard, M. O., Whitwham, A., Keane, T., McCarthy, S. A., Davies, R. M., & Li, H. (2021). Twelve years of SAMtools and BCFtools. GigaScience, 10(2), giab008. 10.1093/gigascience/giab008

Detcharoen, M., Arthofer, W., Steiner, F. M., & Schlick-Steiner, B. C. (2026). Three decades of analyzing *Wolbachia* bacterial endosymbionts in arthropods: Trends and gaps. Parasitology Research, 125(5):159. 10.1007/s00436-026-08694-2

Elbrecht, V., Braukmann, T. W. A., Ivanova, N. V., Prosser, S. W. J., Hajibabaei, M., Wright, M., Zakharov, E. V., Hebert, P. D. N., & Steinke, D. (2019). Validation of COI metabarcoding primers for terrestrial arthropods. PeerJ, 7, e7745. 10.7717/peerj.7745

Elbrecht, V., & Leese, F. (2017). Validation and development of COI metabarcoding primers for freshwater macroinvertebrate bioassessment. Frontiers in Environmental Science, 5:11. 10.3389/fenvs.2017.00011

Forshage, M., & Karlsson, D. (n.d.). The SMTP first-tier sorting. Station Linné. Available at: https://www.stationlinne.se/sv/forskning-research/the-swedish-malaise-trap-project-smtp/taxonomic-units-in-the-smtp/first-tier-sorting/ (accessed June 21, 2026)

Geller, J., Meyer, C., Parker, M., & Hawk, H. (2013). Redesign of PCR primers for mitochondrial cytochrome *c* oxidase subunit I for marine invertebrates and application in all-taxa biotic surveys. Molecular Ecology Resources, 13(5), 851–861. 10.1111/1755-0998.12138

Hague, M. T. J., Shropshire, J. D., Caldwell, C. N., Statz, J. P., Stanek, K. A., Conner, W. R., & Cooper, B. S. (2022). Temperature effects on cellular host-microbe interactions explain continent-wide endosymbiont prevalence. Current Biology, 32(4), 878–888.e8. 10.1016/j.cub.2021.11.065

Hajibabaei, M., Spall, J. L., Shokralla, S., & Van Konynenburg, S. (2012). Assessing biodiversity of a freshwater benthic macroinvertebrate community through non-destructive environmental barcoding of DNA from preservative ethanol. BMC Ecology, 12(1), 28. 10.1186/1472-6785-12-28

Hartop, E., Lee, L., Srivathsan, A., Jones, M., Peña-Aguilera, P., Ovaskainen, O., Roslin, T., & Meier, R. (2024). Resolving biology’s dark matter: Species richness, spatiotemporal distribution, and community composition of a dark taxon. BMC Biology, 22(1), 215. 10.1186/s12915-024-02010-z

Hebert, P. D. N., Cywinska, A., Ball, S. L., & deWaard, J. R. (2003). Biological identifications through DNA barcodes. Proceedings of the Royal Society of London. Series B: Biological Sciences, 270(1512), 313–321. 10.1098/rspb.2002.2218

Hebert, P. D. N., Floyd, R., Jafarpour, S., & Prosser, S. W. J. (2025). Barcode 100K specimens: in a single Nanopore run. Molecular Ecology Resources, 25(1), e14028. 10.1111/1755-0998.14028

Hosokawa, T., Koga, R., Kikuchi, Y., Meng, X.-Y., & Fukatsu, T. (2010). *Wolbachia* as a bacteriocyte-associated nutritional mutualist. Proceedings of the National Academy of Sciences, 107(2), 769–774. 10.1073/pnas.0911476107

Hrček, J., & Godfray, H. C. J. (2015). What do molecular methods bring to host–parasitoid food webs? Trends in Parasitology, 31(1), 30–35. 10.1016/j.pt.2014.10.008

Jalali, S. K., Ojha, R., & Venkatesan, T. (2015). DNA Barcoding for identification of agriculturally important insects. In A. K. Chakravarthy (Ed.), New Horizons in Insect Science: Towards Sustainable Pest Management (pp. 13–23). Springer India. 10.1007/978-81-322-2089-3_2

Johnson, M. G., Gardner, E. M., Liu, Y., Medina, R., Goffinet, B., Shaw, A. J., Zerega, N. J. C., & Wickett, N. J. (2016). HybPiper: Extracting coding sequence and introns for phylogenetics from high-throughput sequencing reads using target enrichment. Applications in Plant Sciences, 4(7), 1600016. 10.3732/apps.1600016

Ju, J.-F., Bing, X.-L., Zhao, D.-S., Guo, Y., Xi, Z., Hoffmann, A. A., Zhang, K.-J., Huang, H.-J., Gong, J.-T., Zhang, X., & Hong, X.-Y. (2020). *Wolbachia* supplement biotin and riboflavin to enhance reproduction in planthoppers. The ISME Journal, 14(3), 676–687. 10.1038/s41396-019-0559-9

Karlsson, D., Hartop, E., Forshage, M., Jaschhof, M., & Ronquist, F. (2020). The Swedish Malaise trap project: A 15 year retrospective on a countrywide insect inventory. Biodiversity Data Journal, 8, e47255. 10.3897/BDJ.8.e47255

Katoh, K., & Standley, D. M. (2013). MAFFT multiple sequence alignment software version 7: improvements in performance and usability. Molecular Biology and Evolution, 30(4), 772–780. 10.1093/molbev/mst010

Kaur, R., Shropshire, J. D., Cross, K. L., Leigh, B., Mansueto, A. J., Stewart, V., Bordenstein, S. R., & Bordenstein, S. R. (2021). Living in the endosymbiotic world of *Wolbachia*: A centennial review. Cell Host & Microbe, 29(6), 879–893. 10.1016/j.chom.2021.03.006

Kolasa, M., Kajtoch, Ł., Michalik, A., Maryańska-Nadachowska, A., & Łukasik, P. (2023). Till evolution do us part: The diversity of symbiotic associations across populations of *Philaenus* spittlebugs. Environmental Microbiology, 25(11), 2431–2446. 10.1111/1462-2920.16473

Leigh, J. W., & Bryant, D. (2015). POPART: Full-feature software for haplotype network construction. Methods in Ecology and Evolution, 6(9), 1110–1116. 10.1111/2041-210X.12410

Letunic, I., & Bork, P. (2007). Interactive Tree Of Life (iTOL): An online tool for phylogenetic tree display and annotation. Bioinformatics, 23(1), 127–128. 10.1093/bioinformatics/btl529

Li, H. (2018). Minimap2: Pairwise alignment for nucleotide sequences. Bioinformatics, 34(18), 3094–3100. 10.1093/bioinformatics/bty191

Linares, M. C., Soto-Calderón, I. D., Lees, D. C., & Anthony, N. M. (2009). High mitochondrial diversity in geographically widespread butterflies of Madagascar: A test of the DNA barcoding approach. Molecular Phylogenetics and Evolution, 50(3), 485–495. 10.1016/j.ympev.2008.11.008

Łukasik, P., & Kolasa, M. R. (2024). With a little help from my friends: The roles of microbial symbionts in insect populations and communities. Philosophical Transactions of the Royal Society B: Biological Sciences, 379(1904), 20230122. 10.1098/rstb.2023.0122

Martin, K. J., & Rygiewicz, P. T. (2005). Fungal-specific PCR primers developed for analysis of the ITS region of environmental DNA extracts. BMC Microbiology, 5(1), 28. 10.1186/1471-2180-5-28

Minh, B. Q., Schmidt, H. A., Chernomor, O., Schrempf, D., Woodhams, M. D., Von Haeseler, A., & Lanfear, R. (2020). IQ-TREE 2: New Models and Efficient Methods for Phylogenetic Inference in the Genomic Era. Molecular Biology and Evolution, 37(5), 1530–1534. 10.1093/molbev/msaa015

Miraldo, A., Sundh, J., Iwaszkiewicz-Eggebrecht, E., Buczek, M., Goodsell, R., Johansson, H., Fisher, B. L., Raharinjanahary, D., Rajoelison, E. T., Ranaivo, C., Randrianandrasana, C., Rafanomezantsoa, J.-J., Manoharan, L., Granqvist, E., Van Dijk, L. J. A., Alberg, L., Åhlén, D., Aspebo, M., Åström, S., … Ronquist, F. (2025). Data of the Insect Biome Atlas: A metabarcoding survey of the terrestrial arthropods of Sweden and Madagascar. Scientific Data, 12(1), 835. 10.1038/s41597-025-05151-0

Nowak, K. H., Hartop, E., Prus-Frankowska, M., Buczek, M., Kolasa, M. R., Roslin, T., Ovaskainen, O., & Łukasik, P. (2025). What lurks in the dark? An innovative framework for studying diverse wild insect microbiota. Microbiome, 13(1), 186. 10.1186/s40168-025-02169-9

Parada, A. E., Needham, D. M., & Fuhrman, J. A. (2016). Every base matters: Assessing small subunit RRNA primers for marine microbiomes with mock communities, time series and global field samples. Environmental Microbiology, 18(5), 1403–1414. 10.1111/1462-2920.13023

Park, E., & Poulin, R. (2020). Widespread Torix *Rickettsia* in New Zealand amphipods and the use of blocking primers to rescue host COI sequences. Scientific Reports, 10(1), 16842. 10.1038/s41598-020-73986-1

Piñol, J., Mir, G., Gomez-Polo, P., & Agustí, N. (2015). Universal and blocking primer mismatches limit the use of high-throughput DNA sequencing for the quantitative metabarcoding of arthropods. Molecular Ecology Resources, 15(4), 819–830. 10.1111/1755-0998.12355

Piñol, J., Senar, M. A., & Symondson, W. O. C. (2019). The choice of universal primers and the characteristics of the species mixture determine when DNA metabarcoding can be quantitative. Molecular Ecology, 28(2), 407–419. 10.1111/mec.14776

Pompanon, F., Deagle, B. E., Symondson, W. O. C., Brown, D. S., Jarman, S. N., & Taberlet, P. (2012). Who is eating what: Diet assessment using next generation sequencing. Molecular Ecology, 21(8), 1931–1950. 10.1111/j.1365-294X.2011.05403.x

Ross, P. A., Ritchie, S. A., Axford, J. K., & Hoffmann, A. A. (2019). Loss of cytoplasmic incompatibility in *Wolbachia*-infected *Aedes aegypti* under field conditions. PLOS Neglected Tropical Diseases, 13(4), e0007357. 10.1371/journal.pntd.0007357

Salter, S. J., Cox, M. J., Turek, E. M., Calus, S. T., Cookson, W. O., Moffatt, M. F., Turner, P., Parkhill, J., Loman, N. J., & Walker, A. W. (2014). Reagent and laboratory contamination can critically impact sequence-based microbiome analyses. BMC Biology, 12(1), 87. 10.1186/s12915-014-0087-z

Scholz, M., Albanese, D., Tuohy, K., Donati, C., Segata, N., & Rota-Stabelli, O. (2020). Large scale genome reconstructions illuminate *Wolbachia* evolution. Nature Communications, 11(1), 5235. 10.1038/s41467-020-19016-0

Seemann, T. (2014). Prokka: Rapid prokaryotic genome annotation. Bioinformatics, 30(14), 2068–2069. 10.1093/bioinformatics/btu153

Shropshire, J. D., Conner, W. R., Vanderpool, D., Hoffmann, A. A., Turelli, M., & Cooper, B. S. (2026). Calibrating and documenting host-switching and evolution of incompatibility loci for two closely related *Wolbachia* clades. *GENETICS*, iyag097. 10.1093/genetics/iyag097

Shropshire, J. D., Leigh, B., & Bordenstein, S. R. (2020). Symbiont-mediated cytoplasmic incompatibility: What have we learned in 50 years? eLife, 9, e61989. 10.7554/eLife.61989

Smith, M. A., Bertrand, C., Crosby, K., Eveleigh, E. S., Fernandez-Triana, J., Fisher, B. L., Gibbs, J., Hajibabaei, M., Hallwachs, W., Hind, K., Hrcek, J., Huang, D.-W., Janda, M., Janzen, D. H., Li, Y., Miller, S. E., Packer, L., Quicke, D., Ratnasingham, S., … Zhou, X. (2012). *Wolbachia* and DNA Barcoding Insects: Patterns, Potential, and Problems. PLoS ONE, 7(5), e36514. 10.1371/journal.pone.0036514

Snell, Q., Walker, P., Posada, D., & Crandall, K. (2002). TCS: Estimating Gene Genealogies.

Song, H., Buhay, J. E., Whiting, M. F., & Crandall, K. A. (2008). Many species in one: DNA barcoding overestimates the number of species when nuclear mitochondrial pseudogenes are coamplified. Proceedings of the National Academy of Sciences, 105(36), 13486–13491. 10.1073/pnas.0803076105

Stahlhut, J. K., Gibbs, J., Sheffield, C. S., Alex Smith, M., & Packer, L. (2012). *Wolbachia* (Rickettsiales) infections and bee (Apoidea) barcoding: A response to Gerth et al. Systematics and Biodiversity, 10(4), 395–401. 10.1080/14772000.2012.753488

Tamura, K., Stecher, G., & Kumar, S. (2021). MEGA11: Molecular evolutionary genetics analysis version 11. Molecular Biology and Evolution, 38(7), 3022–3027. 10.1093/molbev/msab120

Teixeira, L., Ferreira, Á., & Ashburner, M. (2008). The bacterial symbiont *Wolbachia* induces resistance to RNA viral infections in *Drosophila melanogaster*. PLoS Biology, 6(12), e1000002. 10.1371/journal.pbio.1000002

Tourlousse, D. M., Yoshiike, S., Ohashi, A., Matsukura, S., Noda, N., & Sekiguchi, Y. (2016). Synthetic spike-in standards for high-throughput 16S rRNA gene amplicon sequencing. Nucleic Acids Research, gkw984. 10.1093/nar/gkw984

Tournayre, O., Tian, H., Lougheed, D. R., Windle, M. J. S., Lambert, S., Carter, J., Sun, Z., Ridal, J., Wang, Y., Cumming, B. F., Arnott, S. E., & Lougheed, S. C. (2024). Enhancing metabarcoding of freshwater biotic communities: A new online tool for primer selection and exploring data from 14 primer pairs. Environmental DNA, 6(4), e590. 10.1002/edn3.590

Van Der Valk, T., Vezzi, F., Ormestad, M., Dalén, L., & Guschanski, K. (2020). Index hopping on the Illumina HiseqX platform and its consequences for ancient DNA studies. Molecular Ecology Resources, 20(5), 1171–1181. 10.1111/1755-0998.13009

Wagner, D. L., Grames, E. M., Forister, M. L., Berenbaum, M. R., & Stopak, D. (2021). Insect decline in the Anthropocene: Death by a thousand cuts. Proceedings of the National Academy of Sciences, 118(2), e2023989118. 10.1073/pnas.2023989118

Walker, A. W., Martin, J. C., Scott, P., Parkhill, J., Flint, H. J., & Scott, K. P. (2015). 16S rRNA gene-based profiling of the human infant gut microbiota is strongly influenced by sample processing and PCR primer choice. Microbiome, 3(1), 26. 10.1186/s40168-015-0087-4

Weinert, L. A., Araujo-Jnr, E. V., Ahmed, M. Z., & Welch, J. J. (2015). The incidence of bacterial endosymbionts in terrestrial arthropods. Proceedings of the Royal Society B: Biological Sciences, 282(1807), 20150249. 10.1098/rspb.2015.0249

Wickham H (2016). ggplot2: Elegant Graphics for Data Analysis. Springer-Verlag New York. ISBN 978-3-319-24277-4. https://ggplot2.tidyverse.org

Wong, T. K. F., Ly-Trong, N., Ren, H., Demotte, P., Baños, H., Roger, A. J., Susko, E., Bielow, C., De Maio, N., Goldman, N., Hahn, M. W., Dos Reis, M., Vinh, L. S., Huttley, G., Lanfear, R., & Minh, B. Q. (2026). IQ-TREE 3: Phylogenomic inference software using complex evolutionary models. Molecular Biology and Evolution, 43(5), msag117. 10.1093/molbev/msag117

Yu, D. W., Ji, Y., Emerson, B. C., Wang, X., Ye, C., Yang, C., & Ding, Z. (2012). Biodiversity soup: Metabarcoding of arthropods for rapid biodiversity assessment and biomonitoring. Methods in Ecology and Evolution, 3(4), 613–623. 10.1111/j.2041-210X.2012.00198.x

