## Supplementary figures and images for "Insect COI barcoding data as an untapped resource for surveying *Wolbachia* symbioses"

### Supplementary Figure S1

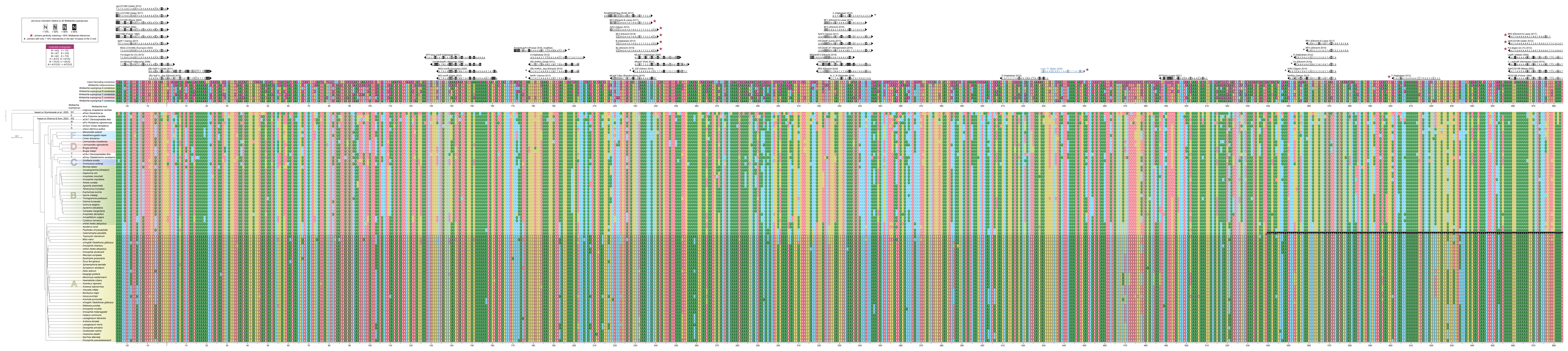
